# Evolution of reproductive strategies in incipient multicellularity

**DOI:** 10.1101/2021.09.13.460035

**Authors:** Yuanxiao Gao, Yuriy Pichugin, Chaitanya S. Gokhale, Arne Traulsen

## Abstract

Multicellular organisms can potentially show a large degree of diversity in reproductive strategies, as they could reproduce offspring with varying sizes and compositions compared to their unicellular ancestors. In reality, only a few of these reproductive strategies are prevalent. To understand why this could be the case, we develop a stage-structured population model to probe the evolutionary growth advantages of reproductive strategies in incipient multicellular organisms. The performance of reproductive strategies is evaluated by the growth rates of corresponding populations. We identify the optimal reproductive strategy, which leads to the largest growth rate for a population. Considering the effects of organism size and cellular interaction, we found that distinct reproductive strategies could perform uniquely or equally well under different conditions. Only binary-splitting reproductive strategies can be uniquely optimal. Our results show that organism size and cellular interaction can play crucial roles in shaping reproductive strategies in nascent multicellularity. Our model sheds light on understanding the mechanism driving the evolution of reproductive strategies in incipient multicellularity. Meanwhile, beyond multicellularity, our results imply a crucial factor in the evolution of reproductive strategies of unicellular species - organism size.

## 1 Introduction

The evolution of multicellularity is viewed as a major evolutionary transition and it has occurred repeatedly across prokaryotes and eukaryotes (Bonner, 1998; Grosberg and Strathmann, 2007; Rokas, 2008; Claessen et al., 2014; Sebe-Pedros et al., 2017; Brunet and King, 2017). With an increase in organism size, phenotypically heterogeneous organisms emerged through cell differentiation (McCarthy and Enquist, 2005; Arendt, 2008; Brunet and King, 2017). Reproductive modes of multicellular organisms may change with organism size and composition. In principle, multicellular organisms could reproduce multiple offspring with distinct cell numbers and organism composition - in contrast to their unicellular ancestors (Michod and Roze, 1999; Ratcliff et al., 2012; Pichugin et al., 2017, 2019; Gao et al., 2019). The number of possible reproductive modes rapidly increases with organism size. For example, for an organism containing three cells, two reproductive strategies are possible: split into three single-celled newborn organisms (1 + 1 + 1) or into a single-celled plus a two-celled newborn organism (2 + 1). For an organism containing ten cells, there are 41 such reproductive strategies, and for a twenty-celled organism, there are 626 reproductive strategies. However, only a few reproductive strategies dominate the tree of life. Some prominent examples abound, such as binary fission producing two single-celled organisms, multiple fission producing many single-celled organisms simultaneously (Suresh et al., 1994; Angert, 2005; Flores and Herrero, 2010), fragmentation reproducing many-celled propagules (Ratcliff et al., 2012) and a special bottleneck reproductive strategy, a multicellular organism producing a single-celled newborn organism repeatedly (Grosberg and Strathmann, 1998; Wolpert and Szathmáry, 2002; Brunet and King, 2017).

The origin and the evolution of reproductive strategies are not well understood. Only a few reproductive strategies have been considered in previous work. The fragmentation mode of producing many-celled propagules has been investigated, in order to understand cell death in yeast (Libby et al., 2014) or to understand the advantages of multicellular life cycles experiencing a unicellular stage (Grosberg and Strathmann, 1998; Michod and Roze, 1999). Previous work has examined the mechanism of life cycle transition from the unicellular stage to the multicellular stage. However, the underlying reproductive strategies are still unknown (Staps et al., 2019). Recent work has also investigated mixed reproductive strategies (Pichugin et al., 2017, 2019), in which the fragmentation mode of an organism is not pre-determined, but selected by natural selection from all fragmentation modes. A subset of reproductive strategies with equal-sized offspring have been investigated in communities with cooperative interactions and deleterious mutations (Henriques et al., 2021). The majority of the literature is focused on the reproductive strategies of homogeneous organisms composed of identical cells. We have recently considered phenotypically heterogeneous organisms (Gao et al., 2019), but cellular interactions were restricted to linear frequency-dependence and we ignored the impact of the organism size. Therefore, it is still unclear how organism size and cellular interaction, together, can shape reproductive strategies.

Organism size confers various advantages to organisms (Kaiser, 2001; Carroll, 2001), such as avoiding predators (Fisher et al., 2016; Kapsetaki and West, 2019), or incentivising the division of labour (Carroll, 2001; Matt and Umen, 2016). Meanwhile, organism size can inhibit growth for different reasons, such as competition for space (Libby et al., 2014) or light (Kapsetaki and West, 2019). Organism size can also affect reproductive strategies as early as nascent multicellularity (Michod, 2007; Solari et al., 2013; Ratcliff et al., 2012; Libby et al., 2014). Field observations are ambiguous about the effects of organism size (Yamamoto and Shiah, 2010; Nielsen, 2006; Li et al., 2014; Wilson et al., 2006; Li and Gao, 2004; Wilson et al., 2010). Here, we consider a broad scope of size effects that can increase, decrease or not change the growth of heterogeneous organisms.

Previous studies have shown that cellular interactions can change reproductive modes (Kaiser, 2001; Solari et al., 2013; Ratcliff et al., 2012). For example, a new phenotype with a higher death rate leads to a reproductive mode of producing propagules among yeast *Saccharomyces cerevisiae* (Ratcliff et al., 2012). Phenotypically heterogeneous organisms could feature diverse cellular interaction forms. Here we study cellular interaction that depends on a minimum threshold of a specific phenotype of an organism. This cellular interaction form has frequently been observed in nature. For example, in response to nitrogen depletion, cyanobacteria differentiate one heterocyst per 10 to 20 vegetative cells (Kumar et al., 2010; Flores and Herrero, 2010). In the genus *Volvox*, along with the germ-soma differentiation (Matt and Umen, 2016), 1 to 20 germ line cells are produced among 500 and 42,000 somatic cells (Shelton et al., 2012).

Thus, both size and composition could affect growth in phenotypically heterogeneous multicellular organisms. We develop a theoretical model to address the evolution of reproductive strategies considering the effects of size and threshold. The size effects could increase or decrease organism growth, while the organism grows fast when its cell number of a phenotype of interest meets a given threshold. Organisms in a population share one common reproductive strategy. Populations differ in reproductive strategies. Reproductive strategies compete with each other via population growth rates. The optimal reproductive strategy maximises the population growth rate. We found that reproductive strategies can co-exist or can dominate others under different conditions. The uniquely optimal reproductive strategy always produces two offspring units.

## 2 Model

We consider multiple populations in which organisms grow and fragment into smaller pieces (see Fig. 1A). The organisms in each population have a unique reproductive strategy. For example, for a population with maturity size *N* = 3, it either has reproductive strategy 1 + 1 + 1 or 2 + 1. In a population with 2+1, mature organisms with three cells produce a single-celled newborn organism and a two-celled newborn organism. The reproductive strategy determines the organism size at which an organism is born and at which size it is mature and reproduces. For the reproductive strategy *n*_1_ + *n*_2_ +… + *n_M_*, newborn organisms have cell number *n_i_* (*i* = 1,…, *M*) and maturity size 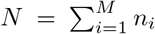. We assume *n*_1_ ≥ *n*_2_ ≥… ≥ *n_M_*. We consider organisms consisting of two cell types: cooperator and defector. This assumption is inspired by the viability investment of organisms for species in the genus *Volvox*, such as *Pandorina, Eudorina*, and *Pleodorina*. At small organism sizes, every cell invests into viability. However, with an increase in the size of the organism some cells gradually decrease their investment into viability (Kirk, 2001, 2005; Matt and Umen, 2016). We refer to the cells contributing to viability as cooperators and the remaining cells as defectors. Newborn organisms may differ in their size and composition in a population. For example, the population with 2 + 1 has five types of newborn organisms: (1,0), (0,1), (2,0), (1,1), and (0, 2), where (*n_D_, n_C_*) shows the number of defectors *n_D_* and cooperators *n_C_*, respectively (see Fig. 1D). Each organism grows incrementally by one cell at a time. During each increment, a cell is selected to divide, and two daughter cells are produced. Each daughter cell can switch to another phenotype independently with a cell-type switching probability, which is *m* = 0.01 in our model. After reaching their maturity size *N*, organisms reproduce via random fragmentation in terms of organism composition. The probabilities of forming different newborn organisms are calculated in Appendix 5.1. The newborn organism follows the same life cycle, growing from newborn to the mature stage, see Fig. 1A.

**Figure 1:**
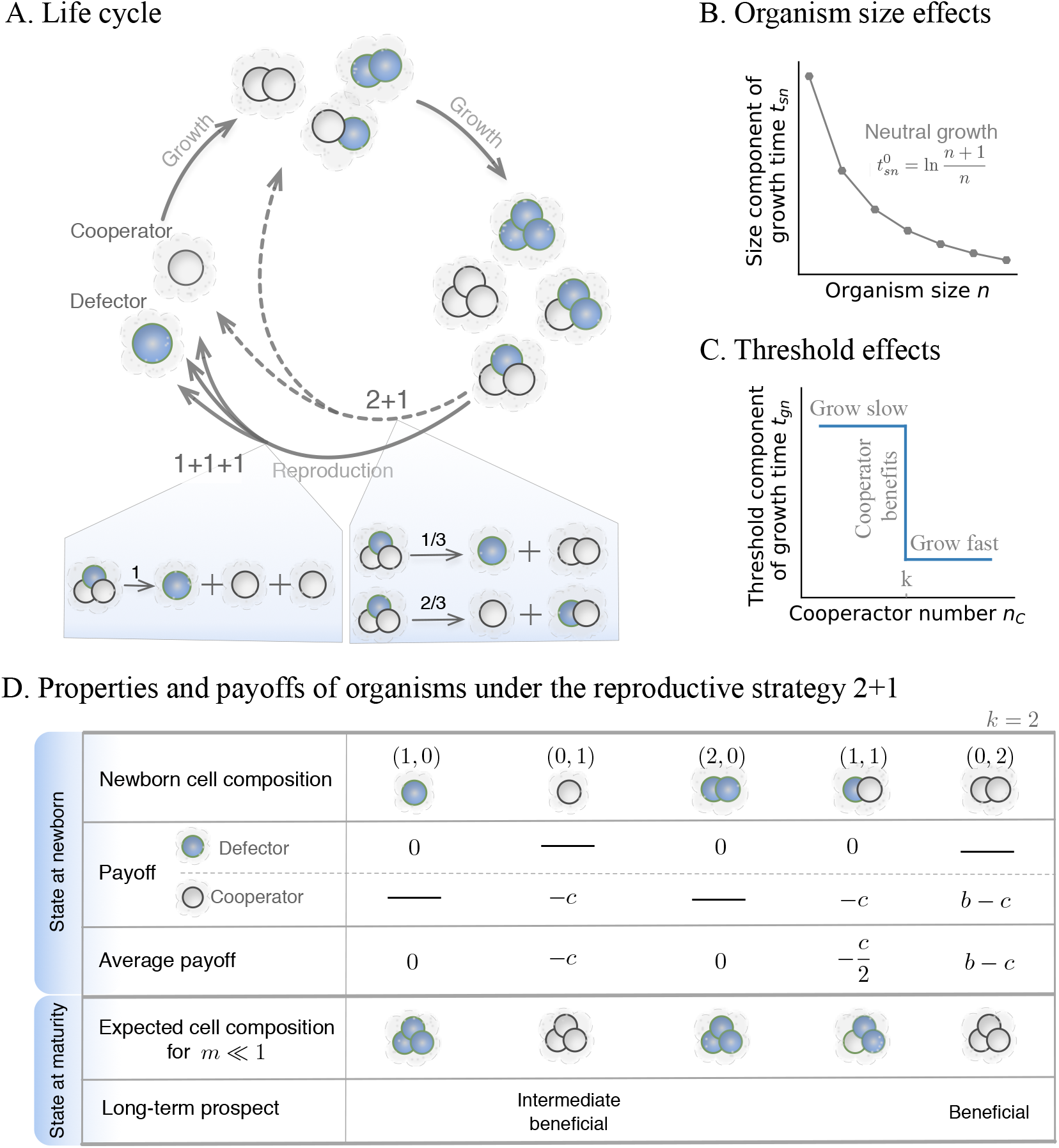
Illustration of a life cycle and the effects of size and threshold. **A.** Example of life cycles with maturity size three. Organisms with different cell compositions at each size stage are illustrated. Two reproductive strategies are shown: 1 + 1 + 1 and 2 + 1. In the shaded area, we show the probabilities of producing different newborn organisms from the mature organism (2,1) under 1 + 1 + 1 and 2 + 1, respectively (see Appendix 5.1 for the calculation). **B.** The organism size *n* affects the growth time of organisms. The grey dots show the neutral condition, where organisms of all sizes have the same growth rate. **C.** Threshold effects on the growth time of organisms. In an organism when the number of cooperators *n_C_* exceeds the contribution threshold *k*, the threshold component of growth time *t_gn_* decreases as in a volunteer dilemma game, see main text. **D.** An example of a population’s newborn organisms and their payoffs under threshold effects. We show the newborn organisms of the population with reproductive strategy 2 + 1. The maturity size *N* = 3. The payoff of each cell in an organism and the average payoffs of organisms are given for *k* = 2. The expected cell composition describes an organism’s cell composition at maturity for *m* ≪ 1. Long-term prospect classifies fast-growing newborn organisms into “beneficial” and “intermediately beneficial”, see main text.

We assume that organisms in populations grow independently without density dependence. Thus, populations follow exponential growth (Tuljapurkar and Caswell, 1997). The population growth rate λ, depending on the number of offspring and the growth time of organisms (De Roos, 2008; Gao et al., 2019), can be calculated as in Appendix 5.2. Since we assume no cell death, the number of offspring of each organism is constant, depending on its reproductive strategy. For example, with the reproductive strategy 2+1, organisms produce two offspring after reproduction. Thus, the population growth rate is determined by the time required for the newborns to mature. We assume that reproduction is instantaneous. The growth time of an organism is then determined by its size and composition as,

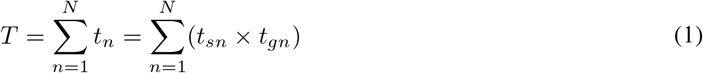

where the *t_n_* is the cell increment time for the organism growing from size *n* to (*n* +1). *t_sn_* and *t_gn_* are the size component and the threshold component of *t_n_*. Next, we discuss how we model *t_sn_* and *t_gn_*.

The size component *t_sn_* depends on the cell number *n* of an organism during growth. Under the neutral condition 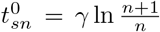, the doubling time of the organism size is independent of the organism size (Gao et al., 2019). Thus, organisms of all sizes have the same growth rate, see Fig. 1B. Without loss of generality, we chose *γ* = 1. To analyze size effects beyond the neutral condition, we screen a large number of values of *t_sn_* around the neutral condition 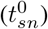, see Fig. 2A. We refer to 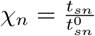 as normalised cell increment components, where *n* = 1,…, *N*. For *χ_n_* = 1, we recover the neutral condition.

**Figure 2:**
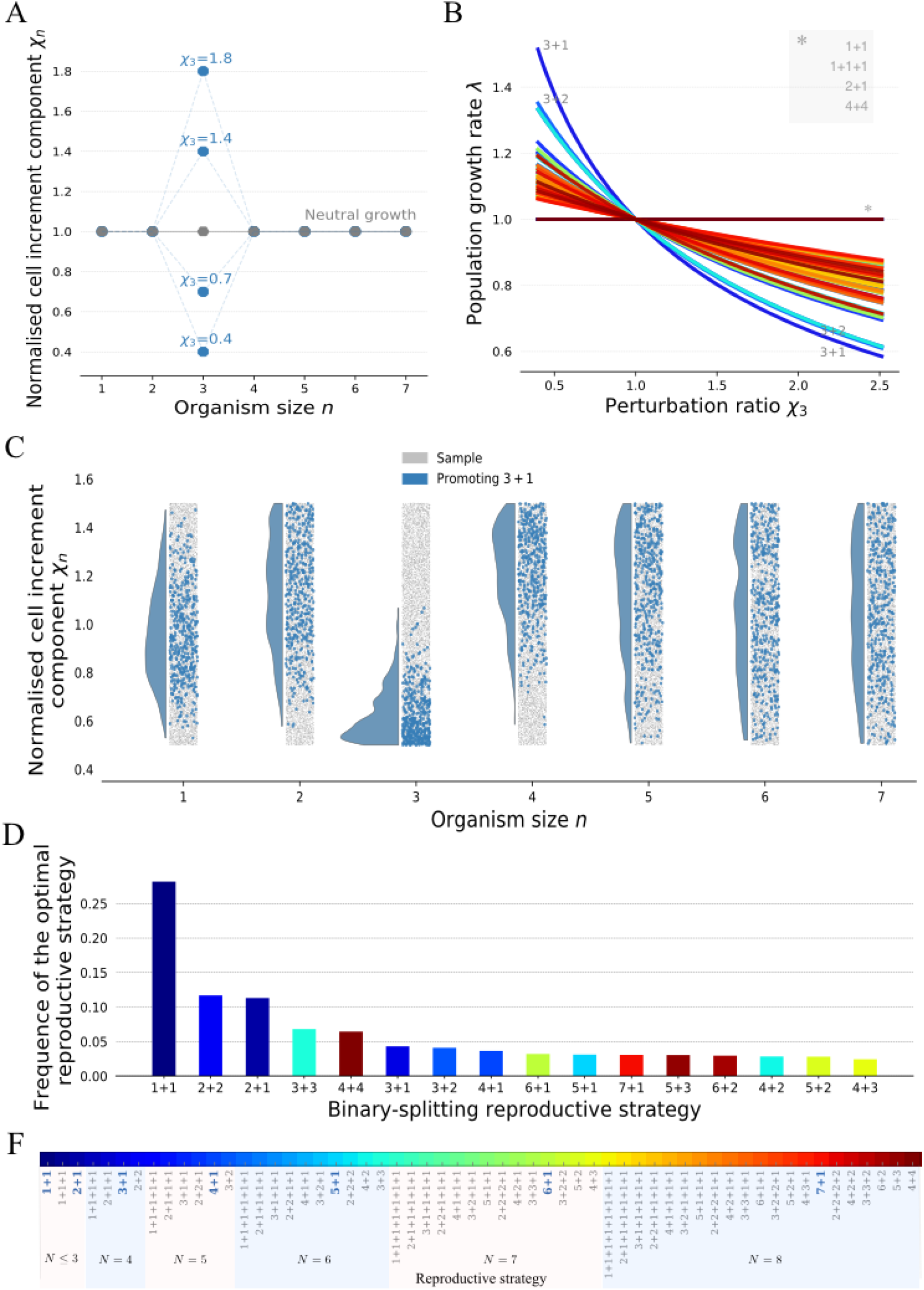
The binary-splitting reproductive strategies are uniquely optimal under the effects of size. **A.** A diagram of perturbations at size *n* = 3. Grey dots are the conditions for neutral population growth *χ_n_* = 1. Blue dots are the perturbed values at size 3 with different strength. **B.** The growth rates of populations with different reproductive strategies under perturbations at size *n* = 3. The asterisk * shows the unaffected reproductive strategies continue to perform equally well. **C.** The distribution of *χ_n_* that promote the reproductive strategy 3 + 1 (in blue) among all samples (in grey). *χ_n_* are drawn randomly from a uniform distribution, where *χ_n_* = 0.5,…, 1.5. A sequence of [*χ*_1_,…, *χ*_7_] is randomly chosen at a time and the optimal reproductive strategy for it is identified. Ten thousand such sequences are investigated in total. **D.** The frequency of observed optimal reproductive strategies under size effects. **F.** The reproductive strategies that have been investigated for the maturity size *N* ≤ 8. The reproductive strategies highlighted in bold blue letters are the optimal ones under a single perturbation *n* = 1,…, 7.

The threshold component *t_gn_* depends on the number of cooperators of an organism. An organism grows faster if the number of its cooperators meets a given threshold *k*, Fig. 1C. There are many methods to construct such compositional threshold effect. Here we choose a volunteer dilemma game (Diekmann, 1985). Consider an organism consisting of *n* cells with *n_D_* defectors and *n_C_* cooperators. When cooperator number *n_C_* meets a contribution threshold *k*, each cell gets a benefit *b*. Each cooperator pays a cost *c* and defectors pay no costs (Fig. 1D),

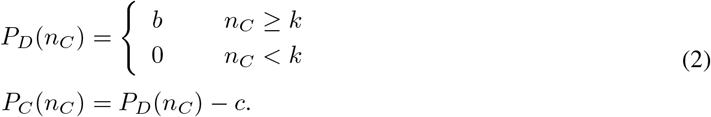

The cell payoffs affect the division probability among these two phenotypes, i.e. which cell is more likely to divide,

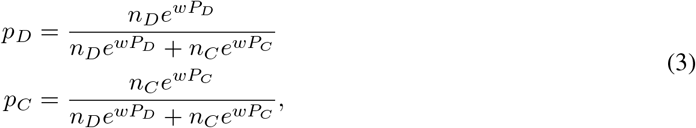

where *p_D_* and *p_C_* are the division probabilities for defectors and cooperators, respectively, and *w* is the intensity of selection (Traulsen et al., 2008). The threshold component *t_gn_* is determined by the payoff *P_D_* and *P_C_*,

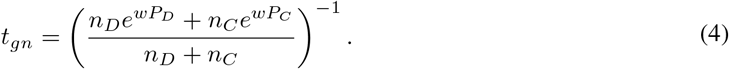

To analyze such threshold effects, we will vary the contribution threshold value *k*.

## 3 Results

### 3.1 The effects of organism sizes on reproductive strategies

To focus on size effects, we assume no threshold effect, *w* = 0. We investigate size effects by perturbing a single normalised cell increment component *χ_n_*, starting from a fully neutral condition *χ_n_* = 1, where *n* = 1,…, 7 (see Fig. 2A). If the organisms of a population are going through a perturbed state at size *n* i.e. *n_M_* ≤ *n* ≤ *N* = ∑ *n_i_*, then its reproductive strategy (*n*_1_ + *n*_2_ +… + *n_M_*) deviates from the neutral condition. Since the population growth rate is inversely proportional to growth time, a perturbation is either advantageous (*χ_n_* < 1, λ > 1) or disadvantageous (*χ_n_* > 1, λ < 1) for population growth. A reproductive strategy is referred to as being promoted (suppressed) when its population growth rate is greater (smaller) than the neutral growth rate 1. A single advantageous perturbation (*χ_n_* < 1) promotes the reproductive strategy of any population with organisms going through the state *n* of the perturbation, i.e. the strategies satisfying *n_M_* ≤ *n* ≤ *N* (Fig. 2B). The performance of reproductive strategies is unaffected when their populations’ organisms do not go through the size under perturbations, i.e. *n* < *n_M_* or *n* > *N*. A single adverse perturbation *χ_n_* > 1 suppresses reproductive strategies that satisfy *n_M_* ≤ *n* ≤ *N*. Among these affected populations, we found that the reproductive strategy *n* + 1 is most affected by perturbations at size *n*. Since the population with reproductive strategy *n* +1 contains *n*-celled newborn organisms, which mature at size *n* + 1, its growth time depends on *χ_n_*. Therefore, under the condition of *χ_n_* < 1 and *χ_k_* = 1 (*k* ≠ *n, k* = 1,…, 7), the reproductive strategy *n* +1 is uniquely optimal. At the same time, the reproductive strategy *n* +1 is most suppressed for *χ_n_* > 1, see Fig. 2B. Analogous to the reproductive strategy *n* +1, the reproductive strategy *n* + 2 is the second most affected reproductive strategy. Similarly, for the rest of reproductive strategies, their population composition determines whether the growth rates are affected or not. The growth rates then determine the performance of reproductive strategies.

When we analyzed general size effects which combine single perturbations at different sizes *n*, we found that the normalised cell increment components determine the optimal reproductive strategies. We observed that the populations of optimal reproductive strategies contain organisms that mostly go through sizes with smaller *χ_n_*. This is illustrated in Fig. 2C and an analytical proof is given in Appendix 5.3 for reproductive strategies with *N* ≤ 3. We found that only the binary-splitting reproductive strategy (producing two offspring) can be uniquely optimal (see Fig. 2D and Appendix 5.4 for the analytical proof). Intuitively, this result is apparent because the fastest-growing newborn organisms in a population with a multiple-splitting reproductive strategy can always be found in another population with a binary-splitting reproductive strategy. For example, the population growth rate of 2+1 + 1 cannot be greater than that of 1 + 1, and 2 + 2 at the same time. Additionally, 1 + 1 is the most frequently observed reproductive strategy in binary-splitting reproductive strategies (see Fig. 2D) because 1 + 1 is the only reproductive strategy that depends on a single cell increment component *χ*_1_. Therefore, for a randomly chosen *χ_n_* (*n* = 1,…, 7), 1 + 1 has a higher probability to be optimal compared to other strategies. Generally, reproductive strategies have lower chances to be optimal when binary-splitting makes organisms go through many cell increment stages.

### 3.2 The effects of thresholds on reproductive strategies

We assume the size effect to be neutral to investigate threshold effects exclusively: *χ_n_* = 1, such that 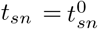. With a threshold at size *k*, newborn organisms of a population with cooperator number *n_C_* ≥ *k* have larger payoffs and thus have shorter growth time, see Eq. (2) and Eq. (4). The growth of different newborn organisms determines the population growth rate. For example, consider all possible newborn organisms in the population with the reproductive strategy 2 + 1: (1,0), (0,1) (2,0), (1,1) and (0, 2), see Fig. 1D. With the contribution threshold *k* = 2, (0, 2) grows fastest as it has two cooperators. (0,1) is the second-fastest-growing newborn organism as it most likely gains benefits by producing a second cooperator during growth. (1,0), (1,1) and (2,0) grow relatively slow because they are less likely to produce at least two cooperators during growth. For convenience, we refer to newborn organisms in a population as “beneficial” if *n_C_* ≥ *k* and “intermediate beneficial” if *n_C_* < *k* and *n_D_* =0. All other newborn organisms are unlikely to reap the benefits of cooperation. The growth rate of a population depends primarily on its beneficial newborn organisms and secondly on its intermediate beneficial newborn organisms. For a low cell-type switching probability, e.g. *m* = 0.01, homogeneous newborn organisms are more abundant than heterogeneous ones. In the long run, we expect that populations mostly contain homogeneous newborn organisms.

For threshold effects, the uniquely optimal reproductive strategies are binary-splitting at the maximum maturity size: 4 + 4, 5 + 3, 6 + 2 and 7 + 1 (see Fig. 3A). The optimal reproductive strategies can be classified into three categories: multiple optima, symmetric binary-splitting 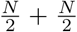 (or 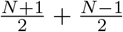) and asymmetric binary-splitting with a *k*-celled newborn organism (*N* - *k*) + *k*. For *k* =1, multiple reproductive strategies are optimal at the same time, see Fig. 3A, B, and C. Since every population contains beneficial newborn organisms, the performances of different reproductive strategies are similar. As *k* increases, the symmetric binary-splitting reproductive strategies 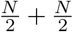 (or 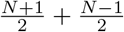) are optimal for 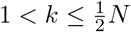, see Fig. 3A B. Newborn organisms with size equal to or greater than *k* have growth advantages, thus intuitively 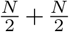 and *k* + (*k* + 1) should have the same performance in population growth. However, we found that only 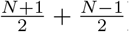 (or 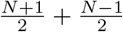) is optimal. The intrinsic composition of the population and the effects of cell-type switching probability *m* = 0.01 determines the results. To understand the growth advantages of the symmetric binary-splitting reproductive strategies with the maximal maturity size, we take 4 + 4 and 3 + 3 at *k* = 3 as an example. For *k* = 3, the population of 4 + 4 contains the beneficial newborn organisms (1, 3) and (0,4). The population of 3 + 3 only contains beneficial newborn organisms (0,3). When a cell-type switching event happens during growth, (0,4) reproduces another beneficial newborn organism (1, 3), while (0, 3) reproduces a non-beneficial newborn organism (1,2). Populations with larger maturity sizes are less affected by the cell-type switching probability as they contain multiple types of beneficial newborn organisms. Finally, when 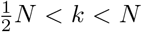, the reproductive strategy (*N* - *k*) + *k* becomes optimal, see Fig. 2A. When 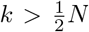, populations can at most have one type of beneficial newborn organism. Next, we explain why the optimal reproductive strategy is (*N* - *k*) + *k* rather than other reproductive strategies such as 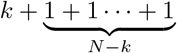 and (*N* − *k* − 1) + *k* + 1. Because of *N* − *k* < *k*, organisms with *N* − *k* cells can only form intermediate beneficial newborn organisms -and only when they are pure cooperators. Larger intermediate beneficial newborns grow faster than smaller ones. We take 3 + 1 + 1 and 3 + 2 under *k* = 3 as an example. 3 + 1 + 1 has the intermediate beneficial newborn organism (0,1) and 3 + 2 has the intermediate beneficial newborn organism (0, 2). During organism growth, (0,1) undergoes two cell increment stages with longer time (larger *t_gn_* due to negative payoffs, see Eq. (4) and Eq. (2)), while (0, 2) only undergoes a single one. Thus, a population with the reproductive strategy 3 + 2 grows faster than one with 3+1 + 1.

**Figure 3:**
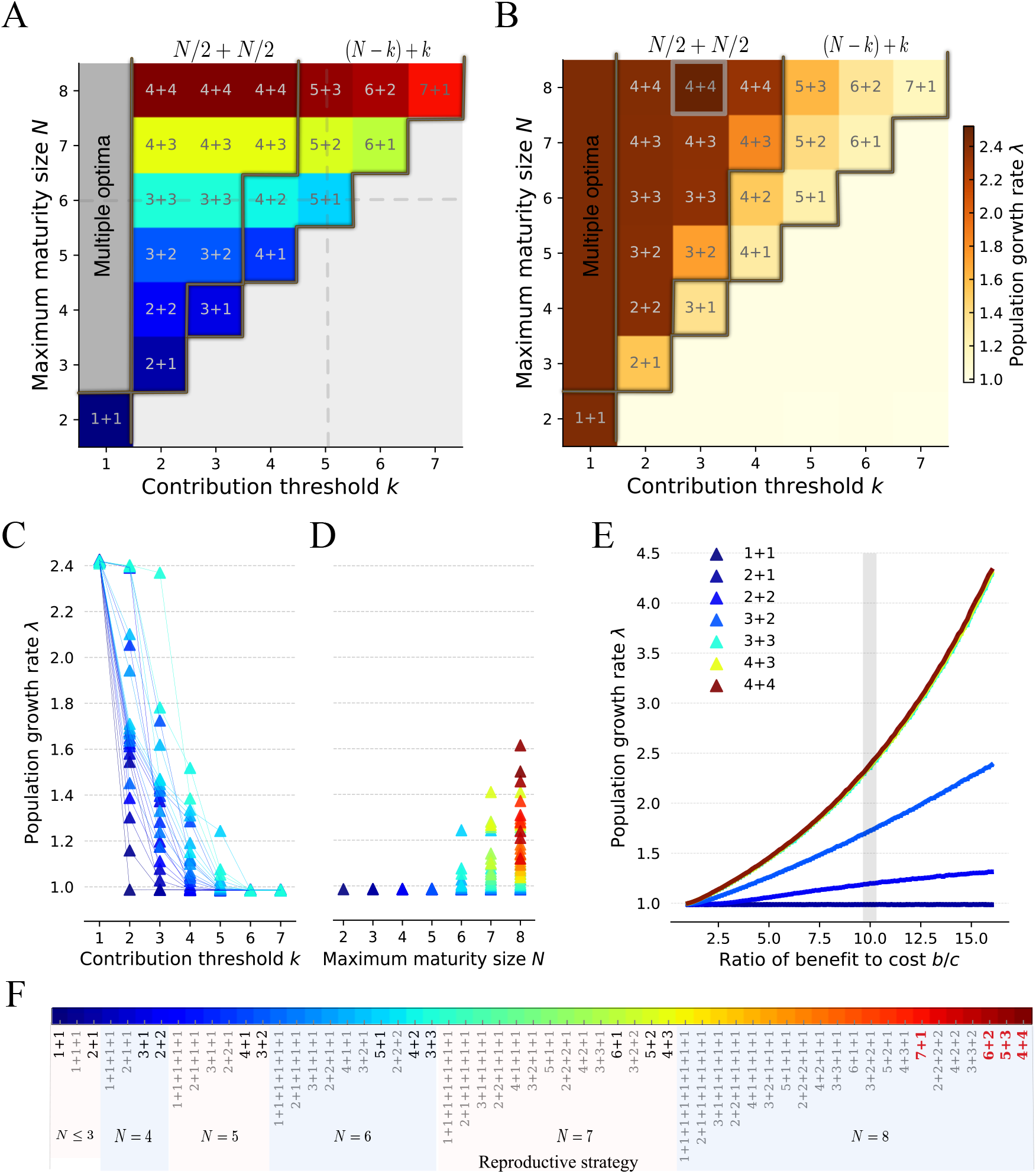
Binary-splitting reproductive strategies are uniquely optimal for threshold effects with *k* > 1. **A.** The optimal reproductive strategies across contribution threshold *k* (*k* < 8) and maturity size *N* (*N* ≤ 8). The dark brown lines (in panels A and B) are the boundaries between multiple optimal reproductive strategies (at *k* = 1), symmetric binary-splitting reproductive strategies, asymmetric binary-splitting reproductive strategies and the section that the threshold never meet. The grey dashed lines indicate the parameter space where we investigated the population growth rate of each reproductive strategy in panel C and D. **B.** The population growth rates of the optimal reproductive strategies in panel A. The highlighted parameter set with *N* = 8 and *k* = 3 is investigated in more detail in panel E. **C.** Population growth rates of different reproductive strategies with *N* ≤ 6 are shown across different contribution threshold *k*. **D.** Population growth rates of different reproductive strategies under contribution threshold *k* = 5 are shown across different maturity size *N* ≤ 8. **E.** The growth rates of populations with symmetric binary-splitting reproductive strategy are shown across to varying ratios of benefit to cost. **F.** The reproductive strategies that have been investigated for *k* ≤ 7 and *N* ≤ 8. The optimal populations that appeared in panel A are highlighted in black. The uniquely optimal reproductive strategies under the threshold effect for *k* =≤ 7 and *N* ≤ 8 are highlighted in bold and red. Parameters for all panels *w* = 0.1, *b* =10, *c* =1 and *m* = 0.01.

Population growth rates decrease with increasing *k*, resulting from reducing the number of beneficial and intermediate beneficial newborn organisms. Especially when *k* ≥ *N*, no reproductive strategies will obtain the benefits of cooperation, and their populations grow slower due to the associated costs, see Fig. 3A, B. Increasing maturity size *N* increases population growth rates of the optimal reproductive strategies because the number of beneficial or intermediate beneficial newborn organisms increases. As expected, population growth rates also increase with the benefit to cost ratio, see Fig. 3B, C, D, and E.

### 3.3 The combined effects of organism sizes and thresholds on reproductive strategies

Finally, we investigate the optimal reproductive strategies under the size and threshold effects combined. For simplicity, we only consider the size effects in the form of a single perturbation. We found that all binary-splitting reproductive strategies *n_i_* + *n_j_* can be uniquely optimal, where *n_i_* and *n_j_* are positive integers, and *n_i_* + *n_j_* ≤ *N* (see Fig. 4A and B). With the combined effects of size and threshold, we found new optimal binary-splitting reproductive strategies that are not optimal either in the effects of single perturbation only or for thresholds only, including 2 + 2, 3 + 2, 4 + 2, 5 + 2, 3 + 3 and 4 + 3. Furthermore, under the beneficial size perturbation, we found *n* +1 (*n* = 1,…, 7) can be optimal both at small and large contribution threshold *k*, see Fig. 4A and B. This is due to the fact that the threshold effects lead to a similar performance of reproductive strategies either at small *k* and at large *k* (Fig. 3B). Therefore, for combined size and threshold effects, the size effects primarily impact the performance of reproductive strategies, see Fig. 4A C. Consequently, the reproductive strategy *n* +1 becomes optimal under an advantageous perturbation, where *n* = 1,…, 7. Newly emerged binary-splitting reproductive strategies have advantages for intermediate contribution thresholds *k*, suggesting that it is an outcome of the trade-off between the effect of size and threshold. For an adverse size perturbation, we found the reproductive strategy *n* + 1 cannot be optimal (Fig. 4B), because the adverse size perturbation leads to poor performance of reproductive strategies that are influenced by the perturbation (see Fig. 2B and Fig. 4D). 7+1 is an exception to this rule, as the threshold effect strongly influence it at *k* = 7. The optimal reproductive strategies observed are those that can obtain growth benefits from threshold effects and avoid the disadvantages from the adverse size effect. For example, 3 + 3 outcompetes 4 + 4 for *k* = 2 when size perturbation occurs at *n* = 7. Both strategies can obtain growth advantages from threshold effects. However, adverse size perturbation decreases the population growth rate of 4 + 4 but has no impact on 3 + 3. Thus the performance of reproductive strategies is the outcome of the trade-off between the effects of size and threshold. Our results suggest that all binary-splitting reproductive strategies can evolve under an appropriate choice of size effects (at a single size) and threshold effects.

**Figure 4:**
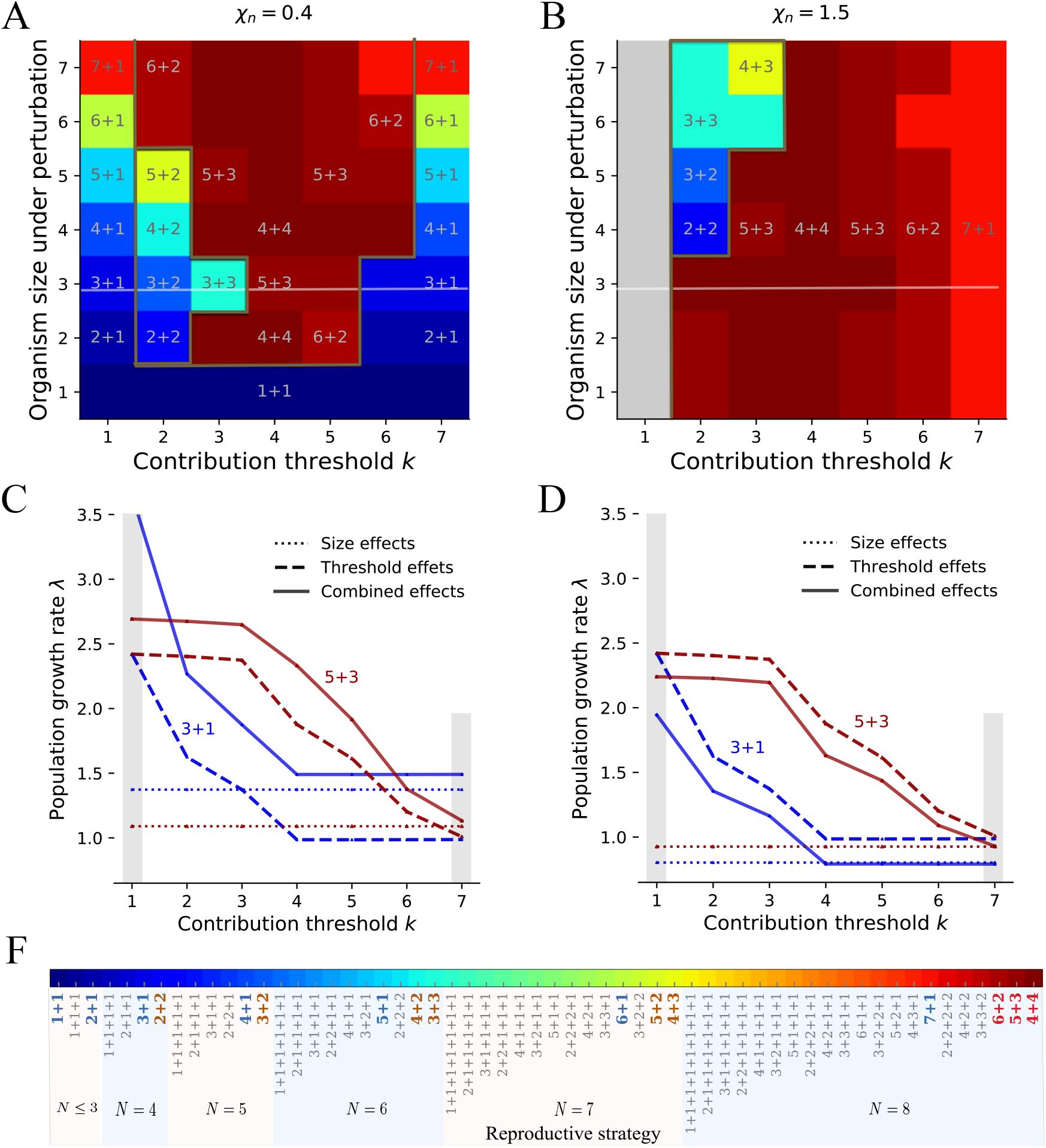
The binary-splitting reproductive strategies are uniquely optimal under the effects of size with a single perturbation and threshold. **A.** Optimal reproductive strategies under the effects of single advantages size perturbations and thresholds. **B.** Optimal reproductive strategies under the effects of single adverse size perturbations and thresholds. In panel A and B, the perturbation only occurs at a single size at a time. The dark brown lines indicate the boundaries of optimal reproductive strategies observed under a single perturbation, threshold effects and both. Note that 7 + 1 is uniquely optimal under either a single perturbation or threshold effects. Reproductive strategies are multiple-optimal under the grey area. The white lines indicate the parameter space where we investigate the population growth rate in panel C and D. **C** and **D.** The population growth rates of reproductive strategies 1 + 3 and 3 + 5 under the effects of a size perturbation at *n* = 3, threshold and both, respectively. In A and C, *χ_n_* = 0.4. In B and D, *χ_n_* = 1.5. **F.** The reproductive strategies that have been investigated for *k* ≤ 7 and *N* ≤ 8. The reproductive strategies in blue are uniquely optimal under the size effect of a single perturbation. The reproductive strategies in red are uniquely optimal under the threshold effects. The reproductive strategies in brown are newly emerged uniquely optimal strategies under both a single perturbation and the threshold effect. Parameters for all panels *w* = 0.1, *b* =10, *c* =1, *m* = 0.01.

## 4 Discussion

Numerous reproductive strategies are conceivable for multicellular organisms, but only recently more attention has been paid to the evolution of reproductive strategies (Tarnita et al., 2013; Pichugin et al., 2017, 2019; Staps et al., 2019; Gao et al., 2019; Pichugin and Traulsen, 2020). Here, we developed a theoretical model considering the effects of size and cell interaction on the evolution of reproductive strategies, impacting organism growth. We considered clonal organisms because of their advantages of purging deleterious mutations and reducing conflicts among cells (Grosberg and Strathmann, 1998, 2007). An alternative way to form multicellular organisms is “coming together”, usually responding to adverse environments (Tarnita et al., 2013; Claessen et al., 2014; Brunet and King, 2017; Amado et al., 2018; Brunet and King, 2017; van Gestel and Wagner, 2021) - but here we entirely focus on “staying together” instead, which typically leads to groups of identical cells when the probability to switch phenotypes is small. We considered cell interaction in the form of a threshold effect, where organism growth depends on the number of cooperators. We sought the optimal reproductive strategy in terms of the largest growth rate of a population. The normalised cell increment component *χ_n_* (*n* = 1,…, *N*) represents the growth time of each cell division. The valve of *χ_n_* and the composition of the population together determine the optimal reproductive strategy. Small *χ_n_* increases the growth rate of reproductive strategies. Contrarily, large *χ_n_* reduces the growth rate of reproductive strategies. We found that only binary-splitting reproductive strategies (producing two offspring) can be uniquely optimal. Specifically, only the binary-splitting reproductive strategy *n* + 1 is optimal under a single size perturbation, where n is the size under perturbation, and *n* = 1,…, 7. Under the threshold effect, the contribution threshold and the cell-type switching probability determine optimal reproductive strategy. We found that the uniquely optimal reproductive strategy is the binary-splitting reproductive strategy with maximum maturity size. We found that all binary-splitting reproductive strategies can be uniquely optimal under the combined effects of size with a single perturbation and threshold. Our results show that only the binary-splitting reproductive strategies can be uniquely optimal. Every binary-splitting reproductive strategy can turn into optimal under the effects of single size perturbation and threshold. Thus, it suggests that they can readily evolve multicellularity under the combined effects of size and threshold.

Our finding that the uniquely optimal reproductive strategies are binary-splitting ones under the size effects coincides with the results in our previous work (Pichugin et al., 2017; Gao et al., 2019). Moreover, we found that the reproductive strategy *n* + 1 with a bottleneck can be uniquely optimal under either size or threshold effects. The result may indicate a new advantage over the previously investigated benefits of decreasing the mutation load and regulating the cell conflict (Grosberg and Strathmann, 1998; Michod and Roze, 1999). Our results also show that multiple reproductive strategies are optimal simultaneously under some special conditions, such as under *k* = 1. This resonates with the observation that one species can possess several reproductive strategies simultaneously in nature (Angert, 2005; Flores and Herrero, 2010; Isaksson et al., 2021; Khanna et al., 2021), such as cyanobacteria, which have reproductive strategies of binary fission, budding and multiple fission. The frequently observed reproductive strategy 1 + 1 among binary-splitting reproductive strategies indicates that 1 + 1 is the best reproductive strategy under uncertain size effects.

In our model, we chose a flexible impact of size on organism growth. Size could have positive, negative or neutral effects on growth at each cell increment. The model assumption is corresponding to studies concerning size effect on growth (Yamamoto and Shiah, 2010; Nielsen, 2006; Li et al., 2014; Wilson et al., 2006; Li and Gao, 2004; Wilson et al., 2010). The form of size perturbations used in our work covers a wide range of size functional forms, including those investigated previously (Pichugin et al., 2017, 2019). We delineated the threshold effect of cellular interactions in a multiplayer volunteer game given the utility of game theory in depicting biological interactions ranging from social foraging to cancer development (Maynard Smith and Price, 1973; Tomlinson, 1997; Dugatkin and Reeve, 2000; Nowak and Sigmund, 2004; Nowak, 2006; Kaveh et al., 2016; Wu et al., 2016; McNamara and Leimar, 2020). We use the volunteer’s dilemma primarily to capture the form of cellular interactions (Diekmann, 1985; Archetti, 2009). Each cell only plays a pure reproductive strategy via its phenotype.

We chose the cell-type switching probability *m* = 0.01, because switching mostly happens under environmental pressure in nature (Gallon, 1992; Claessen et al., 2014). The low switching probability leads to a relatively homogeneous population, which mainly contains homogeneous newborn organisms. If a population has beneficial (or intermediate beneficial) newborn organisms, then homogeneous beneficial (or intermediate beneficial) newborn organisms dominate the population. Although heterogeneous beneficial newborn organisms grow fastest, they are not abundant, because such organisms containing one defector and one cooperator are typically growing into an organism in which there are two defectors.

## 5 Appendix

### 5.1 The probability distribution of newborn organisms

We show the calculation of the probabilities of producing different types of newborn organisms from a mature organism (*n_D_, n_C_*), where *n_D_* + *n_C_* = *N*. The probability to produce the newborn organism type (*n*′*_D_*, *n*′_*C*_) (*n*′_*D*_ + *n*′*_C_* < *N*) is calculated by

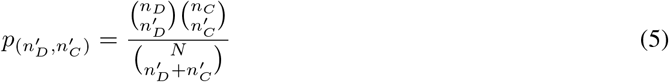

We take the mature organism (1, 2) in a population with reproductive strategy 2 + 1 as an example. The are five newborn organisms: (1,0), (0,1), (2,0), (1,1) and (0,2). The probability of reproducing each newborn organism is shown in Fig. 5.

**Figure 5:**
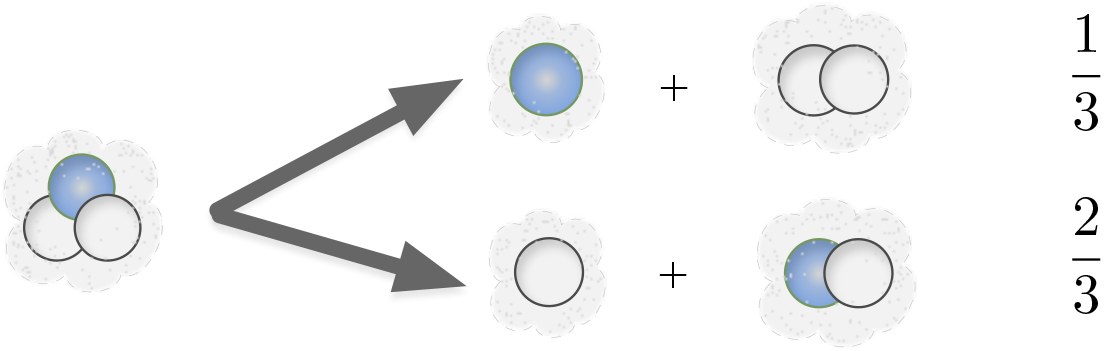
The probability of producing each newborn organism from the mature organism (1, 2) in the population with reproductive strategy 2 + 1. The organism (1, 2) has the probability of 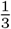 to produce a newborn organism containing one defector and a newborn organism containing two cooperators. It has the probability of 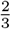 to produce a newborn organism containing one cooperator and a newborn organism containing one cooperator and one defector. However, for small *m* mixed mature groups occur only in small frequency.

### 5.2 Population growth rate

We illustrate the calculation of population growth rates. For the reproductive strategy *n*_1_ + *n*_2_ +… + *n_M_* with maturity size *N*, its population consists of newborn organisms with size *n_i_*, where *i* = 1,…, *M*, 0 < *n_i_* < *N* and 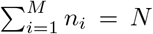. As we consider two cell types, cooperator and defector, an organism with size *n_i_* can have 0, 1,…, *n_i_* cooperators. Therefore, a newborn organism with *n_i_* cells has *n_i_* + 1 possible compositions. We denote the number of newborn organism types of a population by Ω. For example, a population with reproductive strategy 2 + 1 can contain the newborn organisms (1,0), (0,1), (2,0), (1,1) and (0, 2). Here, we would have *N* = 3, *n*_1_ = 1, *n*_2_ =2, *M* = 2 and Ω = 5 (see Fig. 1D). The population growth rate depends on the growth rate of the newborn organisms. We assume that a population contains each type of newborn organisms initially. We track each newborn organism’s growth time and the number of its offspring. We use *T_ij_* to denote the growth time of a *i* type newborn organism until it produces a *j* type newborn organism, where *i, j* = 1,…, Ω. We use *N_ij_* to denote the number of offspring of type *j* offspring produced by the *i* type newborn organism. The growth time *T_ij_* depends on the organism size and the organism composition via Eq. (1). The number of newborn organism *N_ij_* depends on the cell-type switching probability and the cell division probabilities of each cell type. Since organism growth is stochastic, *T_ij_* and *N_ij_* are different for different stochastic trajectories, see (Gao et al., 2019). For example, for the strategy 1 + 1, the newborn organism (0,1) could produce two (1,0), one (1,0) or zero (1,0) with different growth time. To capture the different development trajectories, we simulate the stochastic organism growth and average over *Z* replicates. Then the population growth rate is the largest root of the equation

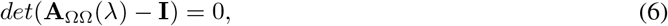

where *A*_ΩΩ_ is a Ω by Ω matrix with elements 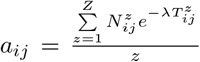 (De Roos, 2008; Gao et al., 2019). Here, 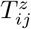 and 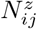 are the growth time and the number of offspring of the newborn organism of size *i* producing an *j* organism in *z*th replication.

The simulation of a population starts with newborn organisms. The newborn organisms differ in their composition, i.e. they have different (*n_D_, n_C_*). For example, for the reproductive strategy 1 + 1, the newborn organisms are of type (1,0) and (0,1). Organisms grow in the following way: In each single step, a cell (cooperator or defector) is selected to divide with its division probability, see Eq. (3). The threshold component of growth time is 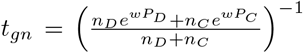 based on Eq. (4). The increment time for the single step is *t_sn_* × *t_gn_*, where we assign values to *t_sn_* according to different scenarios. With the cell division, two daughter cells are produced. Each daughter cell switches to another cell type with a probability *m*. After a single step, we update the number of cooperators and defectors of the organism. Then, the organism repeats the above procedure to grow until reaching its maturity size. Organisms at maturity size produce offspring by random fragmentation. The probability of producing each newborn organism is calculated by Eq. (5) in Appendix 5.1. We obtain the number of offspring produced by the newborn organisms and the growth time (the sum of all time increments) in a single run. We make 5000 replicates of the life cycle of each newborn organism. In the *z*th replication, we record the growth time 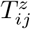 and the number of offspring 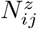 for the *j* type newborn organism producing the *i* type newborn organism. Thus, we have 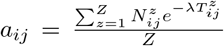, where *Z* = 5000 for our simulations. We numerically recover our analytical results for maturity size *N* ≤ 3, see Appendix 5.3. For *N* ≤ 3, we show that that only the binary-splitting reproductive strategies are uniquely optimal under size effects only in Appendix 5.4. Our remaining conclusions are reached by numerical simulations.

### 5.3 Analytical proof that smaller *χ_n_* determines the optimal reproductive strategy when *N* ≤ 3

For *N* ≤ 3, there are only three reproductive strategies: 1 + 1, 1 + 1 + 1 and 2 + 1. The optimal reproductive strategy is determined by the perturbation with the smaller *χ_n_*. More precisely, the reproductive strategy 1 + 1 is optimal when *χ*_1_ < *χ*_2_ (advantageous perturbation at *n* =1) and 2+1 is optimal when *χ*_1_ > *χ*_2_ (advantageous perturbation at *n* = 2). 1 + 1, 1 + 1 + 1 and 2+1 are optimal when *χ*_1_ = *χ*_2_. The population growth rate of each reproductive strategy is denoted by a subscript. For example, λ_1+1_ describes the population growth rate of the reproductive strategy 1 + 1. The three population growth rates λ_1+1_, λ_1+1+1_, and λ_2+1_ can be calculated by finding the largest eigenvalue of matrix *A* in Eq. (6) in Appendix 5.2. We obtain

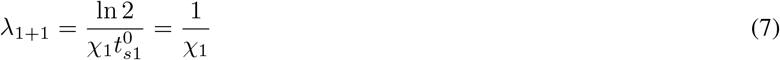

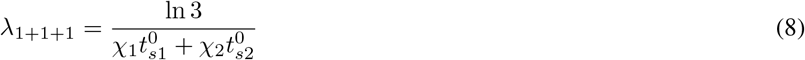

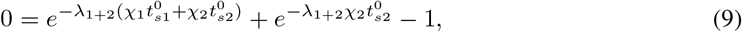

where 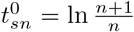 and *n* = 1,2. Eq. (9) only provides an implicit solution for λ_2+1_. The population growth rate is always positive, as there is no cell death in our model setting.

We first focus on *χ*_1_ < *χ*_2_ and prove that the reproductive strategy 1 + 1 leads to faster growth than either 1 + 1 + 1 or 2+1. We start by comparing 1 + 1 with 1 + 1 + 1 for 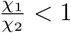,

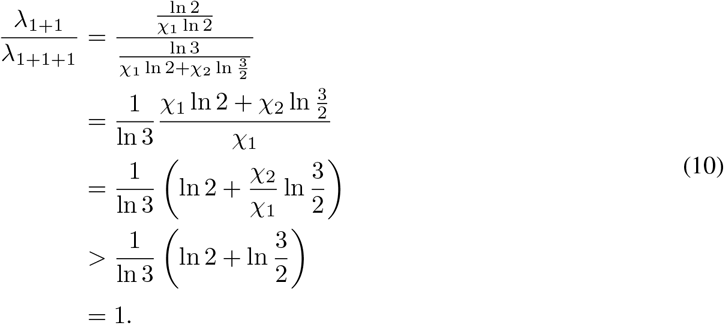

Thus λ_1+1_ > λ_1+1+1_ for *χ*_1_ < *χ*_2_: The reproductive strategy 1 + 1 leads to faster population growth than the reproductive strategy 1 + 1 + 1.

Next we prove that λ_1+1_ > λ_2+1_ for *χ*_1_ < *χ*_2_ by contradiction. If we would have 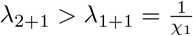, then

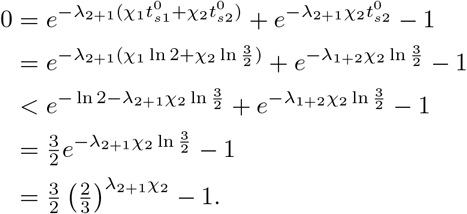

This can be simplified to 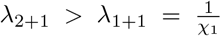 and implies λ_2+1*χ*_2__ < 1 or

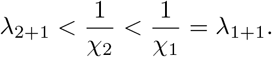

which contradicts the assumption of 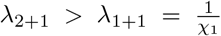. Thus λ_1+1_ > λ_2+1_ for *χ*_1_ < *χ*_2_. Thus the reproductive strategy 1 + 1 is optimal under *χ*_1_ < *χ*_2_.

Now we focus on *χ*_1_ > *χ*_2_ and prove that the reproductive strategy 2 + 1 leads to faster growth than either 1 + 1 or 1 + 1 + 1. We first compare 1 + 1 to 1 + 1 + 1. Since 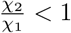, we can revert the argument in Eq. (10) and obtain λ_1+1+1_ > λ_1+1_.

Next we prove – again by contradiction – that λ_2+1_ > λ_1+1+1_ for *χ*_1_ > *χ*_2_. If we would have 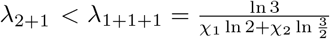, then

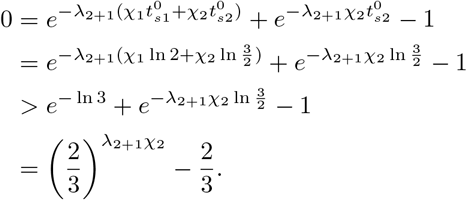

This can be simplified to 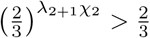 and implies λ_1+2*χ*_2__ > 1 or

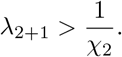

On the other hand, we have for *χ*_1_ > *χ*_2_

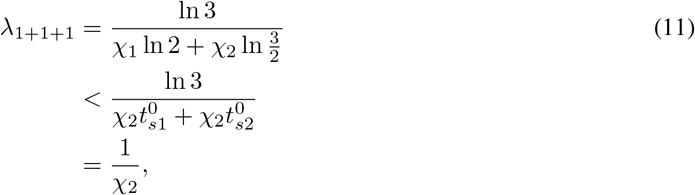

which implies λ_2+1_ > λ_1+1+1_ > λ_1+1_. Thus the reproductive strategy 2 + 1 is optimal for *χ*_1_ > *χ*_2_.

The optimal reproductive strategy under a single size perturbation in the main text is the special case of *χ*_1_ = 1 or *χ*_2_ = 1. Thus, binary-splitting strategies are optimal for *N* ≤ 3. Only for *χ*_1_ = *χ*_2_, all three reproductive strategies of 1 + 1, 1 + 1 + 1 and 2+1 have the same growth rate 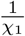. Thus, we have proven that the smaller *χ_n_* determines the optimal strategy. In addition, we found the optimal strategy is either 1 + 1 or 2 + 1, which is consistent with the results that binary-splitting reproductive strategies are optimal under size effects, see Appendix 5.4.

### 5.4 Only the binary-splitting reproductive strategies can be the optimal one under size effects

For size effects only, the number of newborn organism types is reduced as the cell composition does not impact the population growth rate. For example, a population with reproductive strategy 2 + 1 has only two types of newborn organisms: single-celled organisms and two-celled organisms. For the reproductive strategy *n*_1_ + *n*_2_ +… + *n_M_* with 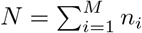, the number of newborn organism types Ω is smaller or qual to *M* (since *n_i_* may be equal to *n_j_*). Therefore, Eq. (6) is reduces to

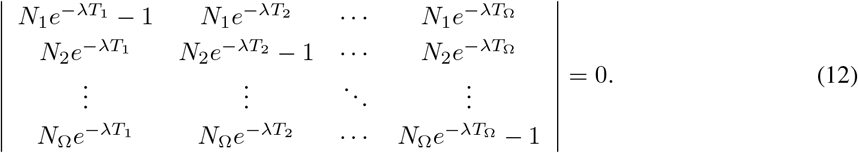

Next, we simplify the determinant on the left hand size of Eq. (12) by changes lines 2 to Ω. We multiply the first row by 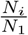 and subtract the result from the ith row, where *i* ∈ [2, Ω]. We obtain

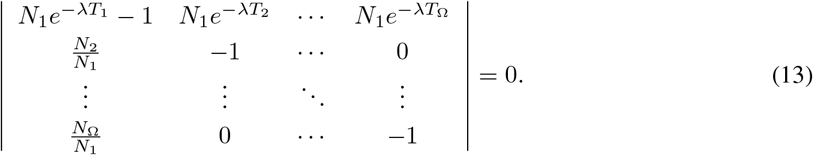

Then we multiply the *i*th column by 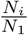 and add it to the first column, where *i* ∈ [2, Ω]. We find

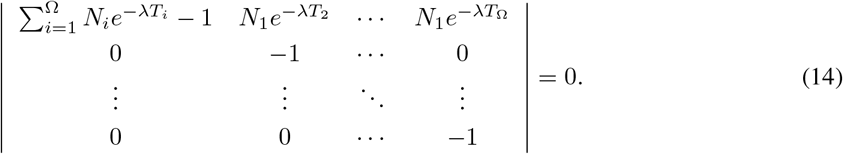

We finally obtain

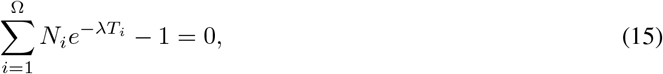

where *i* ∈ [1, Ω]. Since newborn organisms produce identical offspring, *N_i_* is the number of the *i*th type offspring. For example, each organism produces 2 single-celled newborn organisms (the first type) and a two-celled newborn organism (the second type) under 1 + 1 + 2. Thus *N*_1_ = 2 and *N*_2_ = 1. Thus, Eq. (15) can be written in the following equation

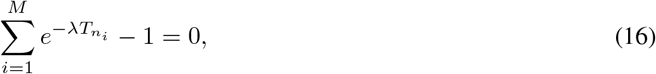

Where *T_n_i__* is the growth time for an organism from newborn size *n_i_* to its maturity size *N*.

To prove that only binary-spitting reproductive strategies can be uniquely optimal, we use a similar method to (Pichugin and Traulsen, 2020). We choose three reproductive strategies *S*_1_ = *n*_1_ + *n*_2_ +… + *n_M_*, *S*_2_ = (*n*_1_ + *n*_2_) +… + *n_M_* and *S*_3_ = *n*_1_ + *n*_2_, where 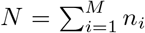. We use λ_1_, λ_2_, and λ_3_ to denote the growth rates of *S*_1_, *S*_2_ and *S*_3_, respectively. The growth rates can be calculated as roots of the equations

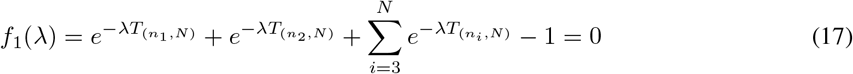

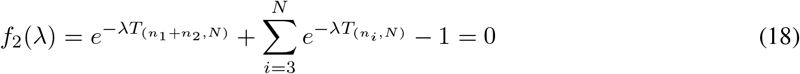

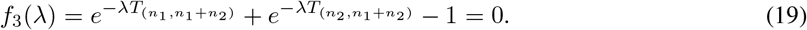

Since the growth time *T* is positive, thus the above equations are monotonically decreasing functions. We multiply Eq. (19) by *e*^-λ*T*_(*n*_1_ + *n*_2_,*N*)_^. Since *T*_(*x,Y*)_ + *T*_(*y,Z*)_ = *T*_(*x,z*)_, we get

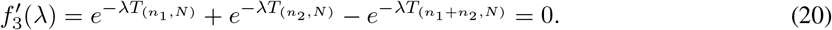

Thus, 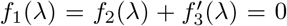. Hence, we have either 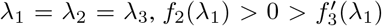 or 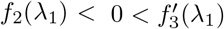 at λ_1_. If *f*_2_(λ_1_) < 0 and 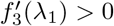, we get λ_2_ < λ_1_ < λ_3_. If *f*_2_(λ_1_) > 0 and 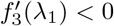, we get λ_3_ < λ_1_ < λ_2_. Thus, uniquely optimal reproductive strategies are always the binary-splitting ones.

## Notes

### Competing Interest Statement

The authors have declared no competing interest.

